# A New Paradigm of Developing Therapeutics to Infectious Diseases by Combining Insights from Nature and Engineering

**DOI:** 10.1101/2020.08.11.246744

**Authors:** Suhyun Kim

## Abstract

Broad host-spectrum antibiotics not only kill pathogens, but also beneficial commensal bacteria of the host microbiome that play crucial roles for health. In Nature, bacteria kill other bacteria much more selectively than antibiotics do. Because there is metabolic cost involved in producing molecules to inhibit others, evolution endowed bacteria with bacteriocins to kill those who are similar enough to compete for the same niche, while leaving more distantly related bacteria, intact. The presence of such narrow host-spectrum antibacterial molecules suggests that by engineering and reprogramming what is found in nature, it may be possible to develop highly effective yet selective therapeutics to infectious diseases, either as purified drugs, or as live bacterial therapeutics. Here, I propose a new paradigm of developing highly selective therapeutics by combining insights from Nature and engineering and applied this against foodborne pathogens, one of the most common causes of bacterial infections for humans.

## Introduction

The gut microbiome consists of various commensal bacteria that play important roles for health, including the production of nutrients and the modulation of the immune system (Belkaid and Hand, 2014; Dodd, et a., 2017). The gut microbiome also affects cognitive functions and emotional states, influences the propensity for obesity, and changes the effectiveness of chemotherapy in cancer patients (Alexander, et al., 2017; Everard, et al., 2013; Sampson, et al., 2016;Yano, et a., 2015). The skin microbiome also serves important functions, including wound healing (Baviera, et al., 2014).

Despite the increased awareness of the importance of the human microbiome, the major treatment option for bacterial infections today is antibiotics. The indiscriminate antibacterial action of broad host-spectrum antibiotics disrupts the host microbiome and prevents it from performing its many functions (Ayres, et al., 2012; Francino, 2016). It also increases the risk of secondary infections by opportunistic pathogens such as *Clostridium difficile* and *Staphylococcus aureus* (Slimings and Riley, 2014; Williams and Gallo, 2017). The central role that the human microbiome plays emphasizes the urgent need to develop new therapeutics that are much more selective than antibiotics. Furthermore, the rise of antibiotic-resistance across the globe also makes it incumbent to develop new solutions to avoid global pandemics caused by resistant bacterial pathogens that do not respond to any currently available antibiotics (Clatworthy, et al., 2007; Cooper and Shlaes, 2011).

In Nature, bacteria kill other bacteria much more selectively than antibiotics do. While killing other competing bacteria is beneficial to procure resources for self in niche competition, there is also metabolic burden involved in producing molecules to kill others (Chao and Levin, 1981; Cza′ra′n, et al., 2001; Riley, 2011). Hence, bacteria are often evolved to produce narrow host-spectrum antimicrobial peptides termed bacteriocins to target those who compete for the same resources, while leaving other, more distantly related bacteria, intact (Riley and Wertz, 2002). Such narrow host-spectrum killing hints that these molecules may be developed into effective yet selective therapeutics to bacterial infections (Riley, et al., 2012). A new paradigm of developing antibacterial therapeutics, then, would be to identify bacteria that are closely related to the target pathogen and use engineering tools and approaches to harness the potential of their native bacteriocin production systems to develop highly selective novel therapeutics (Figure 1A, B).

**Figure 1.**
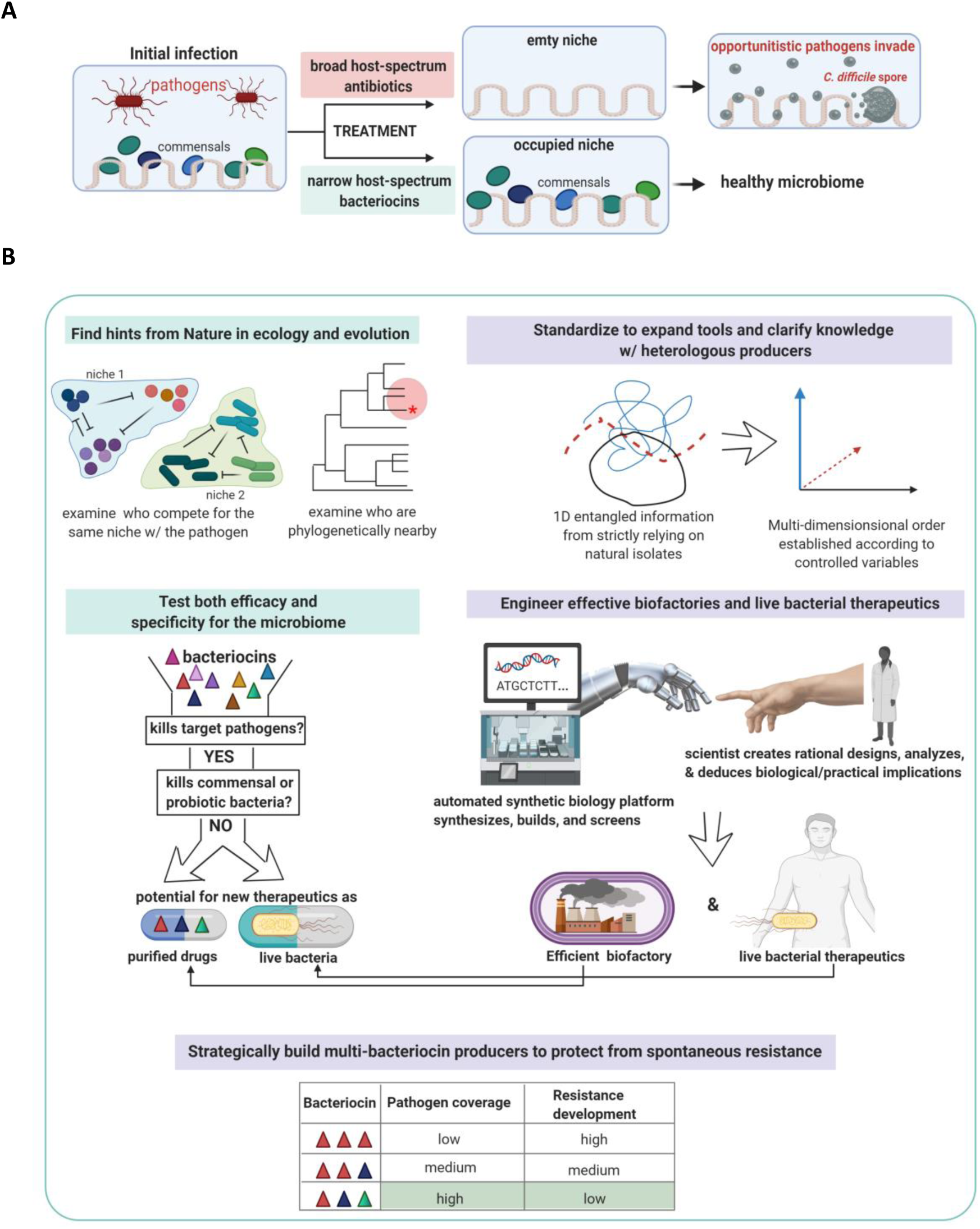
A new paradigm of developing therapeutics to bacterial infections. (A) Effects of broad host-spectrum antibiotics vs narrow host-spectrum bacteriocins on the microbiome (B) A new paradigm of developing therapeutics to infectious diseases proposed and tested in this study.

I applied these principles against food poisoning, one of the most common causes of bacterial infections for humans. One of the main classes of foodborne pathogens are Gram-Negative bacteria including pathogenic *Escherichia coli, Shigella*, and *Salmonella*. In nature, some strains of *E. coli* and *Shigella* naturally produce bacteriocins called colicins (Cascales, et al., 2007; Gratia and Dath, 1925). A study showed *in vivo* efficacy by using purified colicin E1 to kill pathogenic *E. coli* and preventing diarrhea in pigs (Cutler, et al., 2007). Colicin operons have promoters regulated by the bacterial SOS pathway. Normally, LexA proteins bind to the operators and repress transcription. When there is cellular stress or DNA double-strand breaks, which can be experimentally induced by Mitomycin C (MitoC), the SOS pathway activates RecA, which represses LexA and allows transcription (Figure 2A and Cole, et al., 1985; Gillor, et al., 2008; Pugsley, 1987). There are usually three genes in a colicin operon, consisting of the colicin gene that encodes the toxin; the immunity gene that encodes the antitoxin; and the lysis gene to release colicins (Cole, et al., 1985; Pugsley, 1987; Smajs, 1997). Susceptible bacteria have membrane nutrient receptors and porins that are hijacked by these colicins, which disrupt the bacterial cell membranes or act as nucleases to kill the target bacteria (James, et al., 2002; Killmann, 1995; Masi, et al., 1973; Wilmsen, et al., 1990).

**Figure 2.**
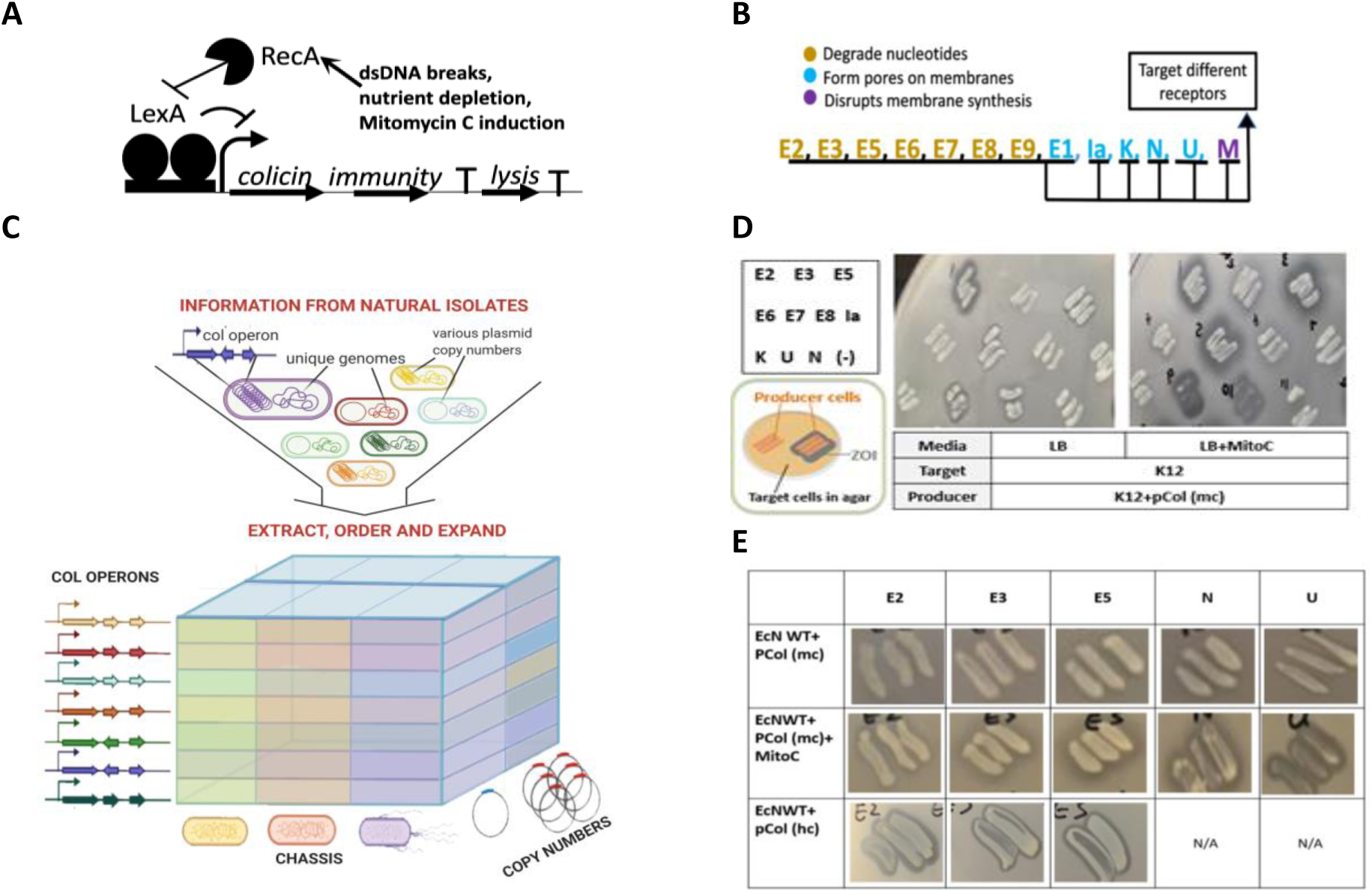
Standardization is key to untangling information from natural isolates and expanding tools and knowledge. (A) A typical colicin operon regulated by the SOS pathway. (B) Colicins used in this study. (C) Standardization using permutations of chassis (host bacteria), copy numbers, and colicin operon sequences. In contrast, effects from these three aspects are not clearly separable when relying on natural isolates. (D) Standardized K12 colicin producers with or without the SOS induction. Each candidate producer is streaked in three lines. ZOI (grey halo) indicates successful killing of the target. (E) Standardized EcN colicin producers with or without the SOS induction and with medium or high copy numbers.

There is substantial barrier in developing bacteriocin-based therapeutics, because bacteriocin operons and gene clusters are often regulated by complex and sensitive systems that are not easily amenable to changes. To overcome current challenges in developing real-life colicin-based therapeutics, I used synthetic biology tools to standardize colicin producers to untangle information from native systems, engineer novel methods to robustly and constitutively produce colicins uninhibited by native control systems, and identify an effective strategy to decrease spontaneous resistance. These advancements and the general approach taken can enable efficient development of bacteriocin-based therapeutics either as drugs in purified forms, or as functionalized probiotics.

## Results

Most studies on colicins utilized naturally isolated colicin producers (Cascales, 2007). Strictly relying on natural isolates makes it impossible to separate the effects from colicin operons, various and often incompletely characterized genomic backgrounds, and plasmid copy numbers. Standardization is key to untangle information from native systems and order it according to relevant variables. Permutations of three different chasses (host bacteria), two different copy numbers, and bare bone colicin operon sequences without unrelated adjacent sequences present in native plasmids were used to build standardized lines (Figure 2C). First, published sequences of colicin operons with different mechanisms of action and target receptors (Figure 2B) were synthesized as DNA parts, cloned into medium copy plasmids (pCol (mc)) and transformed into *E. coli* K12, MG1655 (K12), a model laboratory strain that does not produce any colicins on its own. Modified agar co-culture assay was used throughout the study to test efficacy. In brief, a candidate colicin producer was streaked on top of the agar containing relevant target cells. Effective killing was manifested as zone of inhibition (ZOI)--a halo in the target cell layer surrounding the colicin producer. When the SOS state was induced by the addition of MitoC, K12+pCol (mc) killed wild type K12 (Figure 2D). There was substantial variation among different colicin operons with regards to their inhibitory effects (Figure 2D). Three types of dependency on the SOS induction were observed in the unified genetic and cellular background of K12. The first type strictly required the addition of MitoC (E3, 6, 8). The second type showed the least amount of dependency, creating large ZOI without MitoC (E2). The third type was in-between, creating larger ZOI with Mito C (E5, 7, U, N) (Figure 2D). Previous studies also reported that the induction of the SOS state in native colicin-producing isolates leads to growth defects of the producer cells due to self-lysis from expressing the lysis gene to release colicins (Altieri, et al., 1986; Hakkaart, et al., 1981; Jake and Zinder, 1984; Pugsley and Schwartz, 1983). When tested in a unified, non-native background, however, only colicins U and N—two pore-forming colicins on the list—showed this phenotype. A therapeutically relevant line was generated using a human probiotic *E. coli* Nissle 1917 (EcN) Although EcN is known to encode several bacteriocins, the wild type EcN did not kill K12 or the human foodborne pathogens in LB +/- MitoC. When transformed with pCol (mc), EcN successfully created ZOI on K12 when induced with MitoC (Figure 2E, middle). Unlike K12+pCol (mc), none of the EcN+pCol (mc) strains created ZOI without the full-on induction of the SOS pathway (Figure 2E, top). When transformed with pCol (hc), a high copy number version, EcN created ZOI even without MitoC (Figure 2E, bottom). The third line of standardized producer was generated using *E. coli* NGF for its ability to colonize the mouse gut and allow longitudinal animal studies (Kim, et al., 2018). Like EcN, when transformed with pCol (hc), NGF was able to produce all colicins and create ZOI without the addition of MitoC (Figure S1A, B). Qualitative assessment of the ZOI indicated that K12 and NGF overall lead to stronger production than EcN, suggesting that while EcN is best suited for therapeutic development, it is least suited for *in vitro* overproduction. Overall, results from standardization showed that the strength and the expression control of native colicin operons are significantly influenced by the background genome and copy numbers and that some of the generalized knowledge on colicin biology obtained from studying natural isolates is not as generalizable as it was believed to be.

Using the K12 colicin producers, I tested for the efficacy and the specificity of colicins in the context of the human microbiome. Various strains of pathogenic *E. coli* and *Shigella* all showed susceptibility to colicins, each pathogen with its own susceptibility profile (Table 1). *Salmonella* was not inhibited by any of the tested colicins. Importantly, there was no ZOI on major classes of beneficial commensal and probiotic gut bacteria, including *Bacteroides, Clostridium*, and *Lactobacillus* (Table 1). There was also no inhibition against two major human skin commensal bacteria, *Staphylococcus* and *Cutibacterium* (Table 1).

**Table 1.** Dual-testing of efficacy and specificity using human foodborne pathogens and commensal and probiotic gut and skin bacteria. Parenthesis: source *Colicins in bold indicate stronger effect than other colicins, as judged by larger and/or clearer ZOI **Colicin M was tested using the synthetic operon and was only tested against these two strains.

|  | Species | Strains, serotypes | Colicins that killed |
| --- | --- | --- | --- |
| Foodborne pathogens | EHEC (MGH) | EDL933, annotated as stx - | 2, 3, 5, 6, 7, N |
|  | EPEC (MGH) | E2348/69 | **N\*** |
|  | EAEC (MGH) | *JPN15* | 7, **N** |
|  | EAEC (MGH) | O126:H27 (annotated as *E. coli* 2787) | 7, **U** |
|  | EAEC (ATCC 29552) | Strain CDC 3250-76; serotype o111a,111b:k58:h21; annotated for CVD432 and aggR genes (CVD432+, aggR+) | 7 |
|  | ETEC (MGH) | 10 serotype 029H | **2, 3, 5, 6, 7, N** |
|  | ETEC (ATCC 35401) | Strain H10407; serotype O78:H11; annotated to make LT toxin; Ctx A11; heat stable and heat labile enterotoxins | **2, 3, 5, 6, 7** |
|  | EIEC (MGH) | 15 serotype 029 H | 2, 7, **N** |
|  | EIEC (MGH) | 22 serotype 029H | **2, 3, 5, 6, 7, M\*\*** |
|  | *E. coli* (ATCC 43888) | CDC B6914-MS1; serotype O157:H7 | **2, 3, 5, 6, 7** |
|  | *S. flexneri* (ATCC 12022) | CDC 3591-52 [CIP 104222]; serotype 2b | **U** |
|  | *S. flexneri* (MGH) | M90T; serotype 5a | 2, 3, 5, 6, 7, **U** |
|  | *S. flexneri* (MGH) | 2457T; serotype 2a | **U, M\*\*** |
|  | *S. flexneri* (MGH) | YSH6000T; serotype 2a | **U** |
|  | *S. dysenteriae* (ATCC 9361) | serotype 1 | **N** |
|  | *S. sonnei* (ATCC 25931) | NCDC 1120-66 [CIP 104223] | 2, 7 |
|  | All *Salmonella* strains: *S. typhimurium* (SL1344, from MGH; ATCC 14028; CS401 from MGH; 12023 from MGH; 32-14 14028 from MGH); *S. paratyphiA* (31-10 from MGH; ATCC9150 from MGH); *S. Enteritidis* (30-10 from MGH); *S. typhi* (172-12, LT18, MGH) |  | -<br>(no effect) |
| Probiotic/commensal; gut | *L. plantarum* (BioRhythm) | K1 |  |
|  | *C. butyricum* (Miyarisan) | Miyairi |  |
| Commensal; gut | *B. fragilis* (ATCC 25285) | VPI2553 [EN-2; NCTC 9343] |  |
| Commensal; skin | *C. acnes* (ATCC29399) |  |  |
|  | *S. epidermidis* (ATCC 12228) |  |  |

Efficient and robust overproduction of colicins is a major requirement to develop colicin-based therapeutics, either as purified drugs or as live bacterial therapeutics. One cannot rely on leaky expressions or use MitoC, a carcinogen. However, it has been believed that it is impossible to engineer bacteria that constitutively produce colicins due to the self-lysis aspect of the producer in releasing colicins (Cascales, et al., 2007). Many studies reported excessive self-killing manifested by several log-decrease in the overnight CFU/mL when the expression of the lysis gene was induced (Figure 3A and Altieri, et al., 1986; Cavard, et al., 1989; Mader, et al., 2015; Pugsley and Schwartz, 1983). Another study, in which the colicin E1 plasmid was transformed into a constitutive SOS-state strain called DM1187, reported severe growth defects and corroborated the idea that the whole population of constitutive colicin producers terminate its own existence by irrevocable cell lysis (Figure 3A and Tessman, et al., 1978). As such, the most prominent method currently used in the biotech industry to overproduce colicins for real-life uses involves transgenic plants with microbial DNA—a process that is challenging to scale or adopt and is inherently limited to *in vitro* production (Schulz, et al., 2015).

**Figure 3.**
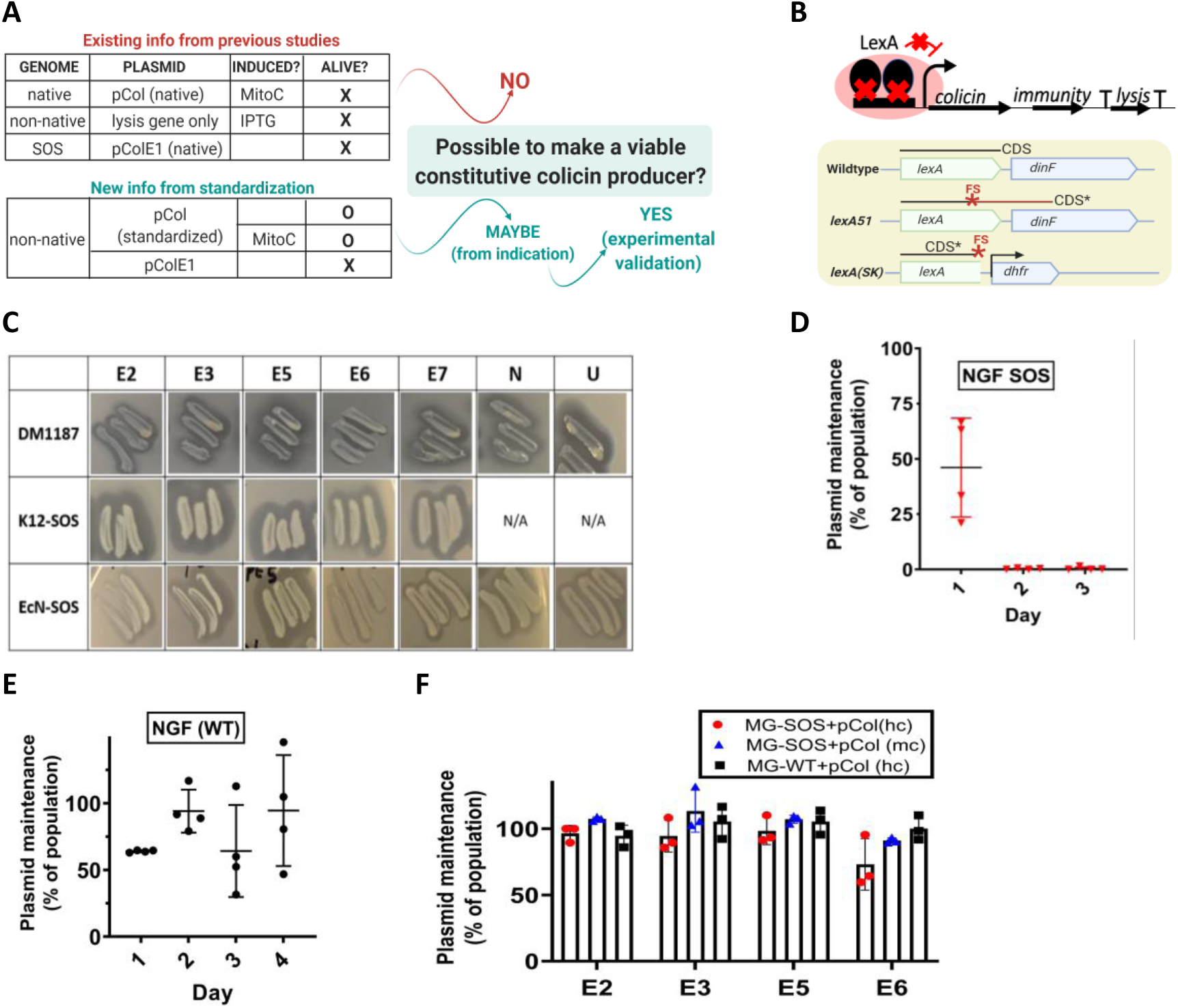
New information from standardization challenged previous beliefs and led to a new hypothesis and successful engineering of robust *in vitro* bio-factories of colicin. (A) Additional knowledge from standardization challenged the previously held beliefs on constitutive production and viability. (B) The *lexA* mutation to genetically establish constitutive SOS state. (C) Successful constitutive production of various colicins. Agar co-culture assay, with K12 in agar in LB. (D-F) Maintenance of standardized colicin plasmids in (D): NGF SOS after passage through the mouse gut; (E): NGF WT after passage through the gut; (F): in MG-SOS and MG-WT after prolonged propagation *in vitro* in non-selective media.

New information gained through standardization suggested that the self-termination of constitutive colicin producers may not be inevitable. First, inducing expressions using MitoC in my standardized colicin producers did not affect growth, except for the pore-forming colicins, N and U (Figure 2D). Second, colicin E1 is also a pore-forming colicin, which may thus have a higher toxicity in production. Finally, when cloning various colicin operons using an automated synthetic biology pipeline (methods), only the colicin E1 operon failed to produce any transformant despite repeated attempts, further suggesting higher self-toxicity. Together, they suggested that the severe growth defects previously reported in overexpressing the lysis genes or the colicin E1 operon may be due more to the imbalance of expressions as well as higher inherent toxicity of colicin E1 compared to other colicins. Accordingly, DM1187 was transformed with seven pCol (mc) plasmids. All of the resulting strains effectively produced colicins (Figure 3C, top). Next, constitutive SOS state was genetically established in K12, EcN, and NGF backgrounds. DM1187 is a natural isolate from an early study on the bacterial SOS pathway and has spontaneously accumulated mutations in the SOS pathway, including *lexA* (as *lexA51*), *recA* (as *recA551ts*; *tif-1*), and *sulA* (as *sfiA11*) (Mount, 1977). As LexA is downstream of RecA in inducing the SOS pathway, I made a loss-of-function mutation in *lexA* (*lexA* (SK)) that results in frameshift and early termination instead of a gain-of-function mutation in *recA*. (Figure 3B). K12-SOS, NGF-SOS, and EcN-SOS all led to successful constitutive production of colicins when transformed with pCol (mc) (Figure 3C, Figure S1C). One exception was EcN+pCol (mc) E6, which only created small ZOI. Strikingly, despite constitutive production, there was no growth defects whatsoever, with the overnight culture CFU/mL of 4×10^9, 2×10^9, and 2×10^9 for K12-SOS, NGF-SOS, and EcN-SOS, respectively. These results show how additional information gained from standardization challenged a previously held belief and led to a new hypothesis that was experimentally validated.

Plasmid loss can easily impede the production of colicins when using engineered bacteria in fermenters or in the gut. Plasmid maintenance during the passage through the mouse gut was examined using NGF-WT or NGF-SOS strains harboring pCol (hc). Fecal samples were plated onto media containing streptomycin to select for all NGF or NGF-SOS cells and onto media containing chloramphenicol to select for only those who maintained plasmids. Unexpectedly, NGF-SOS rapidly lost plasmids, with less than 0.5% of the population maintaining plasmids after two days (Figure 3D). In contrast, the majority of NGF-WT retained plasmids during the four-day post gavage period (Figure 3E). To investigate whether the dramatic plasmid loss also happens *in vitro* when bacteria are propagated in non-selective media, K12-SOS strains with various colicin plasmids were grown overnight or over 2.5 days with 5 subcultures in LB. Most cells retained plasmids at the end (Figure 3F). Therefore, plasmid loss is an unexpected conditional weakness that affects *in vivo* but not *in vitro* applications. Altogether, these results demonstrate that it is possible to engineer extremely robust constitutive colicin producers without the need to add MitoC for induction, antibiotics for plasmid maintenance, or the need to harvest and lyse cells to extract colicins.

The experiment on the *in vivo* stability of plasmids also showed that at least in the wild type genomic background, much of the population could maintain plasmids in the gut for a few days (Figure 3E). Thus, establishing constitutive production in the wild type genomic background would be necessary to develop live bacterial therapeutics for use as functionalized probiotics. Given this, I sought to develop another method to achieve constitutive production without mutating native regulatory pathways. The colicin M operon is a perfect test case, since the native operon for this colicin is dysfunctional as is (Harkness and Olschliiger, 1991). The colicin M operon is a known anomaly among colicin operons, as it lacks a lysis gene. Because of complex regulations and balance involved, there are many potential reasons that a synthetic operon design may fail. These include toxicity from too strong transcription or translation; misbalance between colicins and the immunity proteins; and insufficient release. Finding the balance between optimal production and low burden while avoiding these problems would be difficult and inefficient to achieve with rational design alone. Therefore, after creating multiple designs with various expected expression strengths, I screened/selected against high burden and toxicity by repurposing an automated commercial synthetic biology pipeline, entailing DNA synthesis, cloning, and Next Generation Sequencing Quality Control (NGS QC) (Figure 4A). While the currently proposed value of such automated platforms is delivering sequence-verified plasmids containing the DNA sequences submitted by the customer, failure in this endeavor also provides valuable biological information about one’s DNA designs. I reasoned that while failures in early steps are most likely to be caused by high complexity-DNA structures, failures in later steps, such as the absence of transformants, growth defects, and low scores in NGS QC have biological implications such as high metabolic burden or self-toxicity. Among seven synthetic colicin M operon designs, two passed this automated screening/selection (Figure 4B). These final designs consisted of a synthetic constitutive promoter and synthetic RBSs, the immunity gene and the colicin gene in the reverse order compared to the native colicin M operon, and an ectopic lysis gene from the colicin E2 operon. One of these two designs included the full inter-genic sequences between the 3’ end of the E2 immunity gene and the start of the E2 lysis gene CDS that contain a transcription terminator postulated to prevent self-death by excessive transcription of the lysis gene. The other design differed in that it specifically had this terminator deleted. Functional testing using agar co-culture assay showed that both synthetic operons successfully produced and released colicin M (Figure 4B). Thus, by combining rational design and screening/selection by repurposing an automated synthetic biology pipeline, it was possible not only to build a constitutive system that balances effective production and metabolic burden in the wild type genomic background, but also turn a non-functional colicin operon into a functional one.

**Figure 4.**
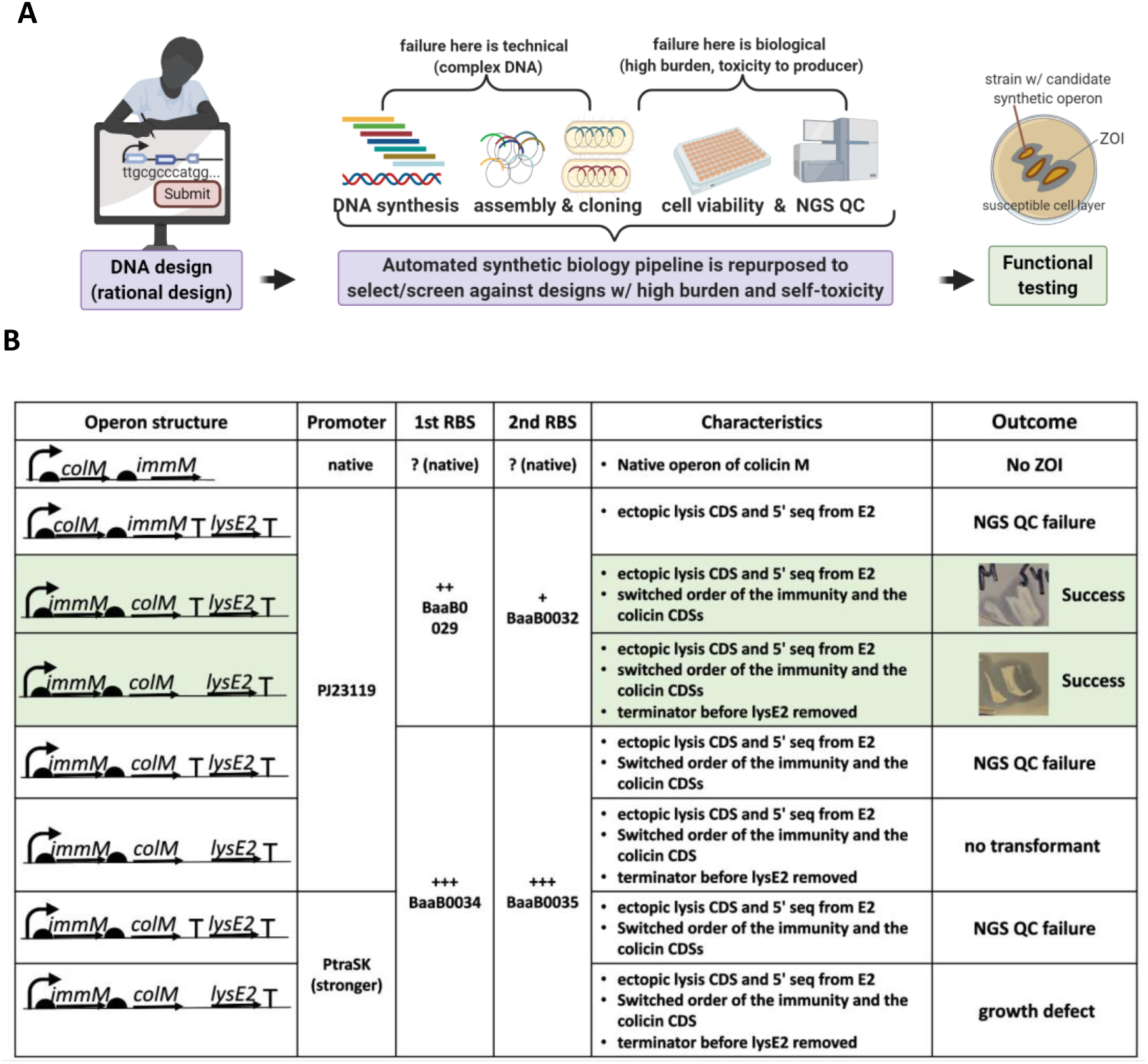
Combining rational design and repurposed synthetic biology pipeline leads to successful engineering of synthetic operons that balance constitutive production and burden. (A) Rational design, screening/selection, and functional testing were combined to build synthetic operons. (B) Seven candidate designs, their main elements, and outcomes. The two shaded in green are successful synthetic colicin M operons that passed the automated screening/selection and the functional testing using agar co-culture assay. +: RBS strengths are relative strengths reported in the iGEM Registry.

Mutations in proteins involved in colicin reception or translocation through the cell membrane are reported to result in resistance to one or more colicins. Experimentally, spontaneously arising colicin-resistant target cells are evidenced by speckles that arise in and around the ZOI created by colicins over time (Calcuttawala, et al., 2015). Whole genome-sequencing of 20 isolates originating from the ZOI of various colicin producers confirmed that these were not contaminating cells and that 18 out of 20 of these resistant isolates had mutations in their known respective main receptors. Thus, the weakest point to spontaneous resistance seemed to be receptors, not translocators or final intracellular targets. Given this observation, I postulated that combining multiple colicins that target different receptors would be a much more effective strategy than combining those that target the same receptor to decrease the occurrence of resistance. To probe this, I engineered the E5/E2 dual-colicin producer that only targets BtuB and the U/M dual-colicin producer that targets both OmpA and FhuA (Figure 5A). The full functionality of these dual-colicin producer strain was confirmed by observing the survival on and the killing of two relevant single-colicin producers. Qualitative and quantitative analysis of spontaneous resistance showed that indeed, combining U/M significantly increased protection against resistance (Figure 5B, C). There is little or no conceivable barrier in combining as many kinds of colicins as possible in concocting drugs using purified colicins. In contrast, in developing live bacterial therapeutics for use as functionalized probiotics, it is essential to maximize the benefit by using only highly complementary colicins, since production of each molecule comes at a metabolic cost to the bacteria. My results suggest that combining colicins that target different receptors is a more effective strategy to reduce the occurrence of resistance development.

**Figure 5.**
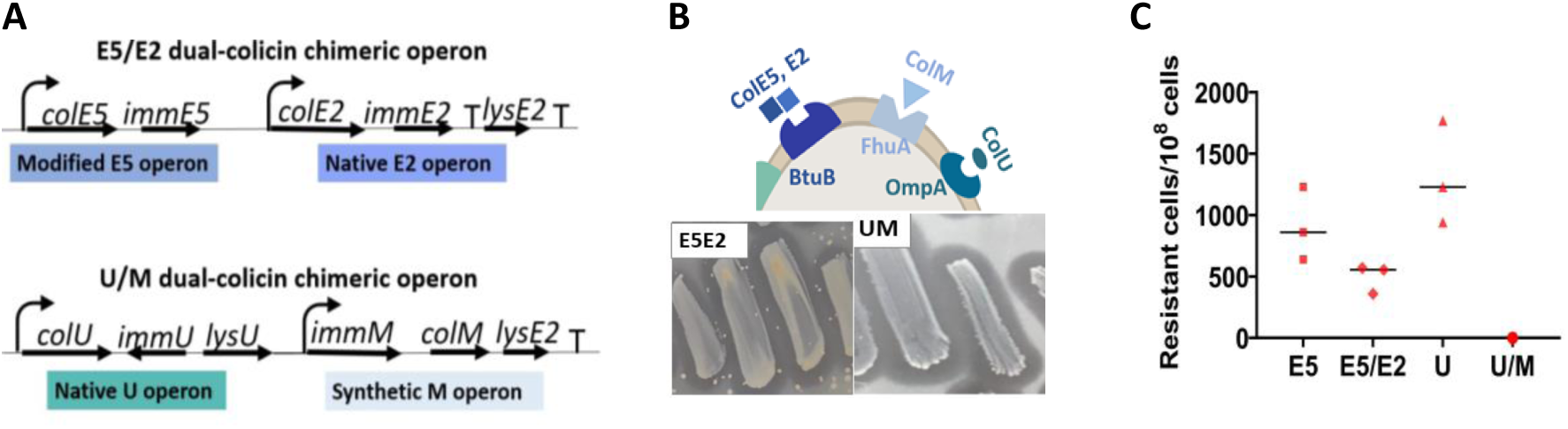
Dual-colicin producers that target different receptors increase protection against spontaneous resistance. (A) Circuit diagrams of two dual-colicin operons. (B) Spontaneous resistance against the E5/E2 and the U/M dual-colicin producers. E5 and E2 target one receptor, whereas U and M target two receptors. Resistant cells in the target layer (K12) become visible as speckles in and around ZOI. (C) The rates of spontaneous resistant cells against single and dual-colicin producers.

## Discussions

In the past 15 years or so, much knowledge has been gained about the crucial roles that the host microbiome plays and the dangers of non-discriminately killing the host commensal bacteria. Despite the increased awareness of the importance of the microbiome, the major treatment option for bacterial infections today is antibiotics, without working alternative therapeutics being available. The first bacteriocin from *E. coli* was discovered in 1925, preceding even the discovery of penicillin (Gratia and Dath, 1925). Although much has been studied on the biology of colicins, the potential of using them as antibacterial therapeutics was largely overshadowed by the subsequent discovery and the widespread acceptance of broad host-spectrum antibiotics. Now is the perfect time to revisit the old, as there is clear understanding of the need to develop more selective antibacterial therapeutics devoid of major problems of antibiotics and there are enabling technologies previously unavailable to aid in this effort. In this study, I proposed an alternative paradigm in developing therapeutics to infectious diseases by harnessing the potential of bacteriocin systems using engineering tools and approaches. I applied these principles against food poisoning as an example.

The lack of susceptibility of beneficial commensal and probiotic bacteria of the human microbiome to colicins is likely due to the divergence of nutrient receptors and porins on the cell membrane that are involved in colicin reception. Although *Salmonella* also did not respond to any of the colicins tested, it further attests to the extreme narrow host-spectrum nature of certain bacteriocins that act more as inter-strain rather than inter-species molecules. Looking for bacteriocins produced by some strains of *Salmonella* may be an effective strategy to identify *Salmonella*-targeting molecules to combine with colicins to build a more complete therapeutic formulation to cover major Gram-Negative foodborne pathogens. Indeed, a recent study reported the discovery of five bacteriocins from *Salmonella* termed salmocins that inhibited 99 major *Salmonella* pathovars (Schneider, et al., 2018). Their specificity should be tested in the context of the human microbiome.

Using synthetic biology tools and approaches, I addressed important challenges and limitations in developing efficient real-life therapeutics based on colicins and colicin-producing bacteria. During standardization, information from natural isolates were untangled, ordered and expanded. Some generalized knowledge about the colicin operon expression regulation, control, and lysis did not hold true when tested in unified, non-native genomic and plasmid backgrounds. This new knowledge obtained from standardization allowed me to engineer constitutive colicin producers as bio-factories or as live bacterial therapeutics. I presented two independent systems to achieve constitutive production of colicins. First, I used genetically established constitutive SOS state in creating robust, constitutive production of colicins without causing growth defects to the producer cells. Second, I created fully functional synthetic operons that can be imported into the wild type genomic background. In doing so, I also generated the first fully functional colicin M producer to my knowledge. The development of both methods was heavily aided by the automation in core synthetic biology technology, which not only reduced the cost and the time of operation, but also allowed me to deduce the metabolic burden and the self-toxicity that my DNA designs inflict on producer cells.

After identifying that receptors are the weakest points to resistance development, I created two different types of dual-colicin producers and showed that combining colicins that target different receptors significantly decrease the occurrence of resistant cells. It remains to be seen whether the reduction in resistance is additive or synergistic when combining multiple colicins. Future studies should also examine how the mutations in colicin receptors affect the pathogenicity of the pathogen. It was reported that the deletion of *ompF* in avian pathogenic *E. coli* decreased the adherence and the virulence of this pathogen (Hejair, et al., 2017). The deletion of *ompA* was also reported to lead to 25- to 50-fold reduction in invasion of *E. coli* into human brain microvascular endothelial cells (Krishnan and Prasadarao, 2012). Thus, when pathogens mutate their receptors to evade colicins, they may also lose part of their pathogenic nature.

In this study, I presented how applying synthetic biology tools and approaches made it possible to harness the potential of nature’s antibacterial strategy towards foodborne pathogens. Given the imminent needs to develop more specific and safer antibacterial therapies than antibiotics, I hope the general approach, the philosophy, and the application of engineering bacteria presented here are widely adopted to develop highly effective and specific antibacterial therapies to many other infectious diseases.

## METHODS

### Bacterial strains

*E. coli* MG1655 (ATCC); DM1187 (CGSG); NGF (Silver lab). For pathogens and commensal and probiotic bacteria, see Table 1.

### Growth media

Non-pathogenic *E. coli* (LB); foodborne pathogens (TSB or GAM); *L. plantarum* (MRS). *C. butyricum* (GAM and anaerobic sachets); *B. fragilis* (pre-conditioned BHI+hematin+cysteine+vitamin K media and anaerobic sachets). *C. acnes* (GAM and anaerobic sachets); *S. epidermidis* (TSB)

### Agar co-culture assay

Agar with appropriate media was autoclaved and cooled to touch. 25mL of the above liquid agar media was transferred into a 50mL falcon tube. Overnight culture of the target cell was diluted to 10^7 CFU/mL and poured onto a petri dish. After solidification, plate was put into a 37’C incubator with the lid slightly open for 30min to allow evaporation of excess moisture.

Afterwards, producer bacteria were gently streaked on top of the agar. The plate was incubated at 37’C overnight. For MitoC induction, final concentration was 0.1ug/mL. To use *B. fragilis* as the target cell, after pouring, plate was dried in an anaerobic jar containing an anaerobic sachet and desiccant to absorb excess moisture for 5h before streaking colicin-producing *E. coli* on top and further incubating for >2d with an anaerobic sachet. Agar containing *C. butyricum* was processed in the same way used for *B. fragilis*. For *C. acnes*, plate was dried with the lid completely open in a hood for 15 min before applying colicin-producers on top and incubating for 2d with an anaerobic sachet. *S. epidermidis* was treated the same way as *E. coli*, except GAM media was used. For *L. Plantarum*, MRS was used to make the agar. After drying, thin layer of LB agar was applied onto this and further dried. Colicin producers were streaked on top of this layer and incubated aerobically.

### Plasmid cloning

pCol (mc) and pCol (mc) dual-colicins were made by using Twist Bioscience’s Cloned Gene Service. They have p15A origin of replication and ampicillin resistance cassette. pCol (hc) was cloned by SK and has a modified pMB1 origin of replication with the *rop* gene deleted. It has chloramphenicol resistance cassette. pCol (hc) was assembled using Golden Gate Assembly-Bsa1, Hifi-v2 (NEB).

### Genetically establishing constitutive SOS state

pKD46 plasmid was used for recombineering. Results were confirmed by colony PCR and sequencing. pKD46 was cured by growing cells at 42’C. For K12 and NGF, *sulA* was also knocked out by replacing the entire *sulA* gene with the EM7 promoter/RBS cassette, followed by *sh ble* for zeocin resistance in the same orientation as *sulA* was originally. For K12, NGF, and EcN, the *lexA (SK)* mutation was created by inserting the EM7 promoter/RBS cassette, followed by *dhfr* for trimethoprim resistance, followed by a transcriptional terminator. This insertion created the following changes in the *lexA* gene and completely removed the *dinF* CDS that originally follows *lexA*. LexA amino acid sequences159-164: KNRAIMS-STOP. Note that during initial selection after cloning and electroporation, cells transformed with a trimethoprim-resistance cassette seems to prefer growth at 30’C for two days and not growth at 37’C.

### Plasmid stability in wild type NGF vs NGF-SOS *in vivo*

Animal protocol was approved by IACUC and ACURO. 8 week-old BalB/C females from Charles River were acclimatized and housed in a BSL2 animal facility. Mice were administered with approximately 10^10 CFU consisting of equal mixture of NGF or NGF-SOS with pCol (hc) E2, E3, E5, or E6. Feces were weighed, homogenized in PBS in a continuous vortex, centrifuged for 2 min at 100g to pellet down only large particles but not the bacteria, diluted in PBS and plated onto relevant selection plates using a spiral plater (Eddy Jet2, Neutec).

### Synthetic operons

PJ23119 and RBS sequences are from the iGEM Registry of Standard Biological Parts. PtraSK is a strong, synthetic promoter that was originally devised for an unrelated project, in which traR-binding sequence (ACGTGCAGATCTGCACA) was inserted between the –35 and –10 of the pJ23119 promoter. This promoter had demonstrated much stronger expression than PJ23119. The DNA sequence of the terminator within the ectopic E2 sequence that was specifically deleted in some synthetic operon designs is: TGGCATTCTTTCACAACAAGGAGTCGTT. Twist BioScience’s Cloned Gene Service for automated synthesis, cloning, growing, and NGS QC was repurposed into a screening/selection tool.

### Dual-colicin operons

The E5/E2 dual-colicin operon was made by attaching the two operons in series, but with the lysis gene from E5 removed. The U/M operon was made by attaching the native U operon and a synthetic M operon in series.

### Resistance rates

Since agar media contained 10^7 CFU/mL of target cells, it was possible to calculate how many bacteria were contained in the relevant volume of ZOI from which speckles were counted, by multiplying the area of ZOI estimated by using ImageJ with the height of the agar. The rate of resistance was calculated by dividing the number of speckles by the total number of cells in that ZOI.

### Software

Geneious, Bowtie and Mauve (DNA designs, assembly); BioRender (figure diagrams); GraphPad Prism (plots); ImageJ (ZOI area and speckle counting)

### DNA acquisition numbers and DNA sequences

(to be deposited upon paper acceptance)

## Author contribution

SK: all aspects of investigation—conceptualization, design, experimentation, analysis, and manuscript writing

## Acknowledgement

Dr. Goldberg at MGH for sharing foodborne pathogens. Dr. Silver and Dr. Way at Harvard Medical School for fund-raising (DARPA-Biological Control).

---

**Figure S1.**
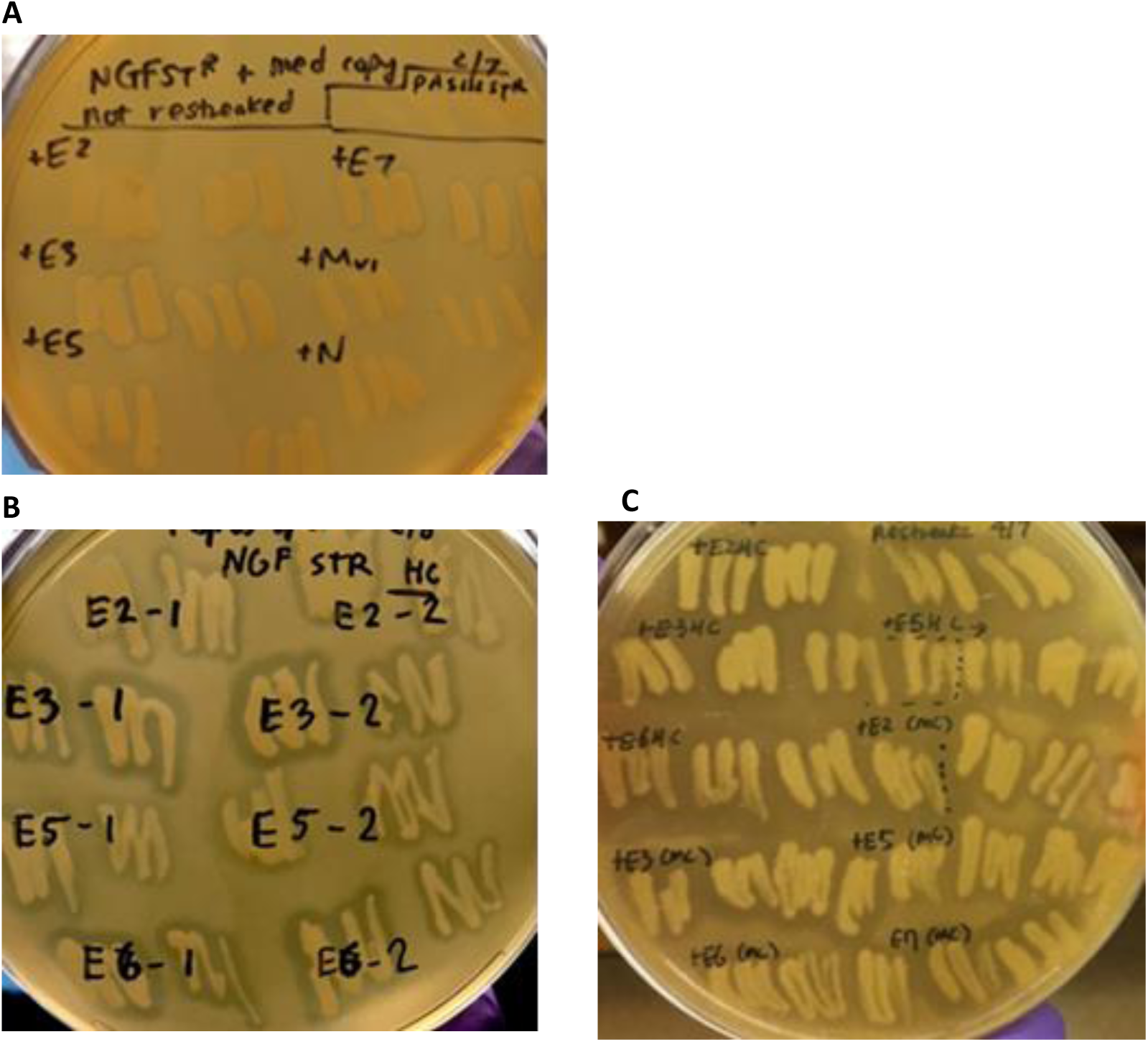
Characterization of NGF-based colicin producers. (A) Wild type NGF with pCol (mc). Only NGF+pCol (mc) E3 and E7 lead to small ZOI without the MitoC induction. (B) Increasing the plasmid copy number leads to successful production even without MitoC, as is manifested by NGF+pCol (hc) E2, E3, E5, and E6 creating clear ZOI. Numbers after dash simply indicate replicates. (C) Genetically established constitutive SOS state resulted in obvious and substantial increase in the uninduced production. NGF-SOS+pCol (hc) E2, E3, E5, and E6 (first four sets) lead to much larger ZOI compared to the wild type with high copy plasmids in (B). Even with medium copy plasmids, NGF-SOS lead to the creation of clear ZOI, in contrast to the wild type NGF+pCol (mc) in (A). Media used here is GAM media without MitoC. Pathogen embedded in the agar as the target cell is pathogenic *E. coli* EIEC 22 serotype 029 H.

